# Auditory tract maturation in early infancy: Links to speech production

**DOI:** 10.1101/2024.12.31.630971

**Authors:** Feihong Liu, Yaoxuan Wang, Jinchen Gu, Jiameng Liu, Xinyi Cai, Han Zhang, Jun Feng, Zhaoyan Wang, Hao Wu, Dinggang Shen

**Author notes:** Contributing authors. Feihong Liu and Yaoxuan Wang contributed equally to this work.

## Abstract

Infants exhibit remarkable language acquisition abilities, supported by highly plastic neural substrates that dynamically interact with early speech experiences. However, the developmental mechanisms of these neural substrates and their specific role in speech acquisition remain incompletely understood. Here, we present NeoAudi Tract (NAT), a robust automated toolbox for extracting the full set of auditory tracts in infants from birth to 24 months using **3**T diffusion MRI data. By characterizing the microstructural changes in these tracts, we demonstrate a gradual and continuous maturation process of the auditory system. Additionally, we identify significant correlations between auditory tract maturation and both *fine-motor skills* and *expressive language **t***-scores from the Mullen Scales of Early Learning tests. Our findings highlight the role of the auditory system in speech production and indicate the intertwined development of auditory and motor systems that underlies speech acquisition, particularly during perceptual reorganization.

## 1 Introduction

Infants are unparalleled language learners, endowed with highly plastic neural substrates that rapidly adapt to early speech experiences, enabling the swift acquisition of essential communicative and cognitive skills [1–3]. On the one hand, ongoing neural development progressively enhances the ability to detect and recognize speech patterns, from basic linguistic structures to complex, high-level ones [4, 5]; on the other hand, these speech experiences shape the maturation of the underlying neural circuits [6]. A hallmark of speech acquisition during this period is the shift from a universal sensitivity to phonetic contrasts across all languages in early infancy, to a more specialized sensitivity attuned to the language(s) infants regularly hear by the end of the first year. This transformation is commonly referred to as *perceptual reorganization*, *perceptual attunement*, or *perceptual narrowing* [7–10]. Crucially, the maturation of the relevant neural substrates in this period not only facilitates communicative speech abilities [11], but also lays a foundation for broader intellectual skills, including reading, mathematics, and general cognitive ability (*e.g.*, IQ) in later childhood [12–16]. Despite this growing body of evidence, the precise mechanisms by which these neural substrates develop to support perceptual reorganization for speech remain poorly understood. Clarifying how early speech experience interacts with the neural circuits may therefore be pivotal for advancing our understanding of both language development and broader cognitive outcomes.

Recent research indicates that speech processing relies on the functional integration of the auditory and motor systems [17–20]. While auditory and motor processing predominantly contribute to speech perception and production, respectively [21–24], they also interact in each other’s processing [18, 25]. The auditory system, for instance, may engage in motor learning [26], thereby influencing speech production [27]. However, these mechanisms remain less understood than the motor system’s role in speech perception [28].

Numerous studies have demonstrated the critical role of the motor system in speech perception [29–34], showing that speech perception is an active process rather than a passive process [35, 36]. Notably, the motor system compensates for speech perception in noisy or distorted cases by enhancing the activities of components in the motor system [37, 38]. Direct stimulation of articulatory regions in the motor cortex specifically interferes with the recognition of corresponding phonemes [18, 39, 40], and restricting articulatory movements of the lips or tongue similarly disrupts phoneme identification [41–44]. Taken together, these findings underscore the critical importance of linking aural (speech perception) and oral (speech production) processes [45]. Although associations between auditory and motor systems have been observed in infants [46], as young as 3 months of age [47], *they may not be present at birth* [48]. Consequently, current research increasingly focuses on early-life sensorimotor experiences that facilitate the establishment of these sensory-motor associations [49–52], rather than on isolated sensory or motor experiences [19, 53–55].

Early-life experiences drive plastic changes in neural substrates, particularly in white matter, thereby enabling efficient transmission and synchronization of neural firing across functionally segregated brain regions [6, 56, 57]. The formation of a mature, interconnected brain network serves as the basis for a range of cognitive functions, including speech [58, 59]. Thus, understanding the dynamics of white matter maturation can provide critical insights into the neural mechanisms underlying speech acquisition and cognitive development [60, 61]. However, despite increasing evidence on the importance of white matter maturation, the specific contributions of auditory tracts to speech production, especially in the context of perceptual reorganization, remain unclear, underscoring the need for further investigation.

To non-invasively investigate the development of the white matter tracts in infants, diffusion MRI (dMRI) has emerged as the primary and widely employed technique [62–67]. Yet, extracting auditory tracts in infants poses unique challenges, due to their smaller size, complex structures, and considerable spatiotemporal variability of developing tissues [68, 69].

Early investigations into auditory tract development have relied predominantly on histological observations, which provide a rough developmental timeline [70–72]. With subsequent advances in neuroimaging, these anatomical insights have been significantly enriched through non-invasive characterization of the tiny and complex auditory nuclei [73, 74]. Given the minute size and complex structures of these brainstem nuclei and tracts, high-resolution imaging is required. Recent methodologies in adults have employed either *ex vivo* 3T MRI [75] or *in vivo* imaging with 7T scanners [74, 76]. For infants, additional challenges arise. Shorter scanning times are required to accommodate unsettled conditions, which results in lower data quality [77]. Moreover, smaller brain sizes combined with spatiotemporal asynchrony lead to high variability in data distribution, complicating data acquisition, processing, and analysis [69, 78]. Although BabyAFQ leverages the infant atlas developed in our prior studies [79] and a classical registration framework to extract 24 major tracts [78], it remains incapable of extracting auditory tracts. To address this gap, we constructed an auditory tract atlas by combining tract density imaging (TDI) with structural MRI (sMRI) data [68]. Despite these advancements, tools for extracting auditory tracts in infants remain scarce, highlighting the continued need for developing more specialized, automated approaches.

In this study, we propose an automated toolbox, *NeoAudi Tract* (NAT), for extracting the full set of auditory tracts spanning peripheral, subcortical, and cortical regions in infants from birth to 24 months. Leveraging 3T MRI data from the most comprehensive longitudinal Baby Connectome Project (BCP) dataset [80], we characterize the developmental trajectories of these auditory tracts, revealing a gradual and continuous maturation process. Auditory tracts closer to peripheral regions exhibit higher maturity, while those nearer to cortical regions show lower maturity. Notably, significant correlations were identified between auditory tract maturation and *fine-motor skills* and *expressive language t*-scores from the Mullen Scales of Early Learning (MSEL) tests, suggesting a potential influence of auditory processing on speech production. Furthermore, our results indicate that during early infancy, auditory tract development is intricately linked with the motor system, raising the possibility that the auditory system may, via motor mediation, play a role in attuning phonetic sensitivity during the first year of life. However, the precise maturation mechanisms through which the auditory system integrates with the motor system to subserve speech acquisition, particularly in the context of perceptual reorganization, remain unclear, highlighting the need for further investigation into the complex interplay between sensory and motor systems.

## 2 Results

Using our proposed NeoAudi Tract (NAT), we extracted the full set of auditory tracts from the longitudinal multi-shell high angular resolution diffusion imaging (HARDI) data in BCP [80] — one of the largest cohorts of infants with both high-quality neuroimaging data and behavioral scores assessed by the MSEL [81]. Our analysis focused on charting the developmental trajectories of these auditory tracts from birth to 24 months and examining their correlations with *t*-scores in four MSEL domains: *visual reception*, *fine-motor skills*, *receptive language*, and *expressive language*.

Of the entire dataset, 246 data samples (164 infants) passed the quality control (QC) procedures, as detailed in the Method section. Table 1 provides the sample sizes by age month. For correlation analysis, we selected MSEL *t*-scores from these data samples that were acquired within a 2-month window of their dMRI scanning time points, ensuring close temporal alignment. Fig. 1 illustrates the distribution of matched subjects across these intervals (122 infants, 189 data samples), including a subset older than 24 months to fully utilize the available longitudinal data.

**Table 1.**
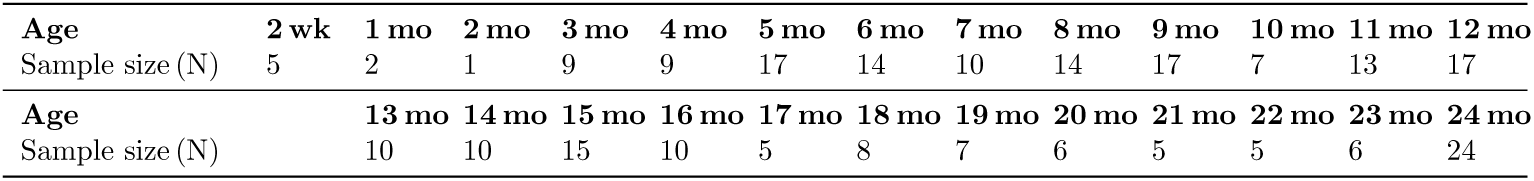
Number of data samples per age month after quality control.

| Age | 2 wk | 1 mo | 2 mo | 3 mo | 4 mo | 5 mo | 6 mo | 7 mo | 8 mo | 9 mo | 10 mo | 11 mo | 12 mo |
| --- | --- | --- | --- | --- | --- | --- | --- | --- | --- | --- | --- | --- | --- |
| Sample size (N) | 5 | 2 | 1 | 9 | 9 | 17 | 14 | 10 | 14 | 17 | 7 | 13 | 17 |
| Age | 13 mo | 14 mo | 15 mo | 16 mo | 17 mo | 18 mo | 19 mo | 20 mo | 21 mo | 22 mo | 23 mo | 24 mo |  |
| Sample size (N) | 10 | 10 | 15 | 10 | 5 | 8 | 7 | 6 | 5 | 5 | 6 | 24 |  |

**Fig. 1.**
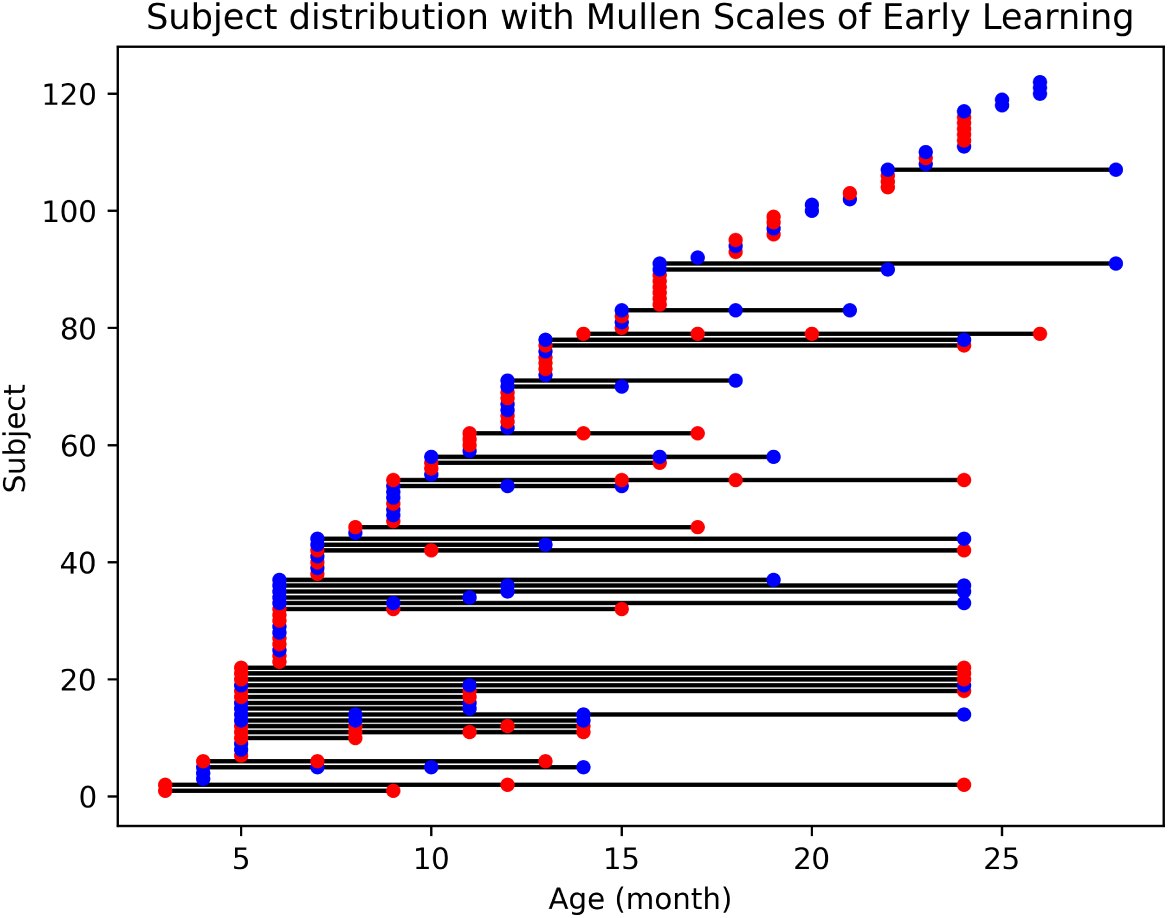
Distribution of subjects in the longitudinal study with paired MSEL scores and diffusion MRI data across different ages. The study includes data from 122 subjects (63 females and 59 males). Blue points represent males, red points represent females, and horizontal black lines connect data samples from the same individual.

### 2.1 Depiction of extracted auditory tracts

Fig. 2 illustrates the extracted auditory tracts and the aligned ROIs at six representative ages (3, 6, 9, 12, 18, and 24 months). These tracts and ROIs span peripheral, subcortical, and cortical regions [68, 82]. More specifically, in the peripheral ear, the cochlea (CO) captures and converts speech vibrations into electrical pulses, which are transmitted via the cochlear nerve to the cochlear nucleus (CN). The signals then sequentially reach other brainstem and thalamic nuclei in the subcortical regions [82, 83] before ultimately arriving at the primary auditory cortex (PAC) in the superior temporal cortex [84]. In total, we extracted 14 auditory tracts in this study, reflecting the comprehensive coverage of peripheral, subcortical, and cortical auditory pathways. As illustrated in Fig. 2, all auditory tracts were successfully identified, demonstrating the effectiveness of NAT for infant auditory tract extraction. From birth to 24 months, the streamlines of each tract become progressively denser, corresponding to the steady increase in brain size and myelination [69, 85]. In younger infants (*e.g.*, 3 months), the extracted tracts appear noticeably smaller and sparser, reflecting the relatively underdeveloped brain structure at this early stage. By contrast, at 24 months, the tracts are visibly larger and denser, mirroring the ongoing maturation of white matter [86].

**Fig. 2.**
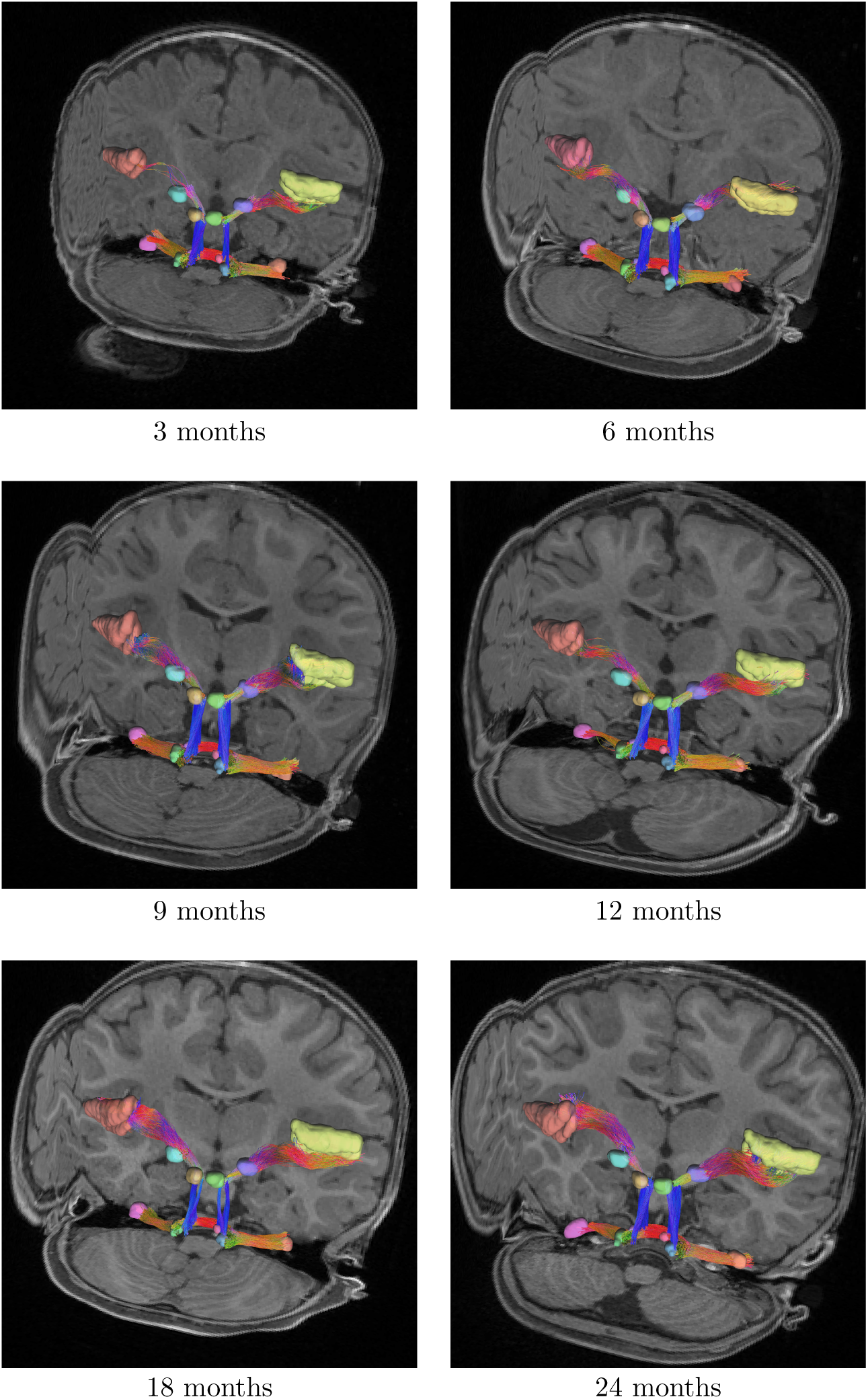
Extracted auditory tracts and aligned ROIs across six age periods (3, 6, 9, 12, 18, and 24 months). Tract size and streamline density increase progressively with age, reflecting ongoing brain development during early infancy.

To evaluate the effectiveness of NAT, we assessed the alignment of auditory nuclei — initial and terminal regions of these tracts — which determines the reliability of tract extraction. As illustrated in Fig. 3, we compared three commonly used registration strategies for aligning adult templates with infant sMRI data:

**Fig. 3.**
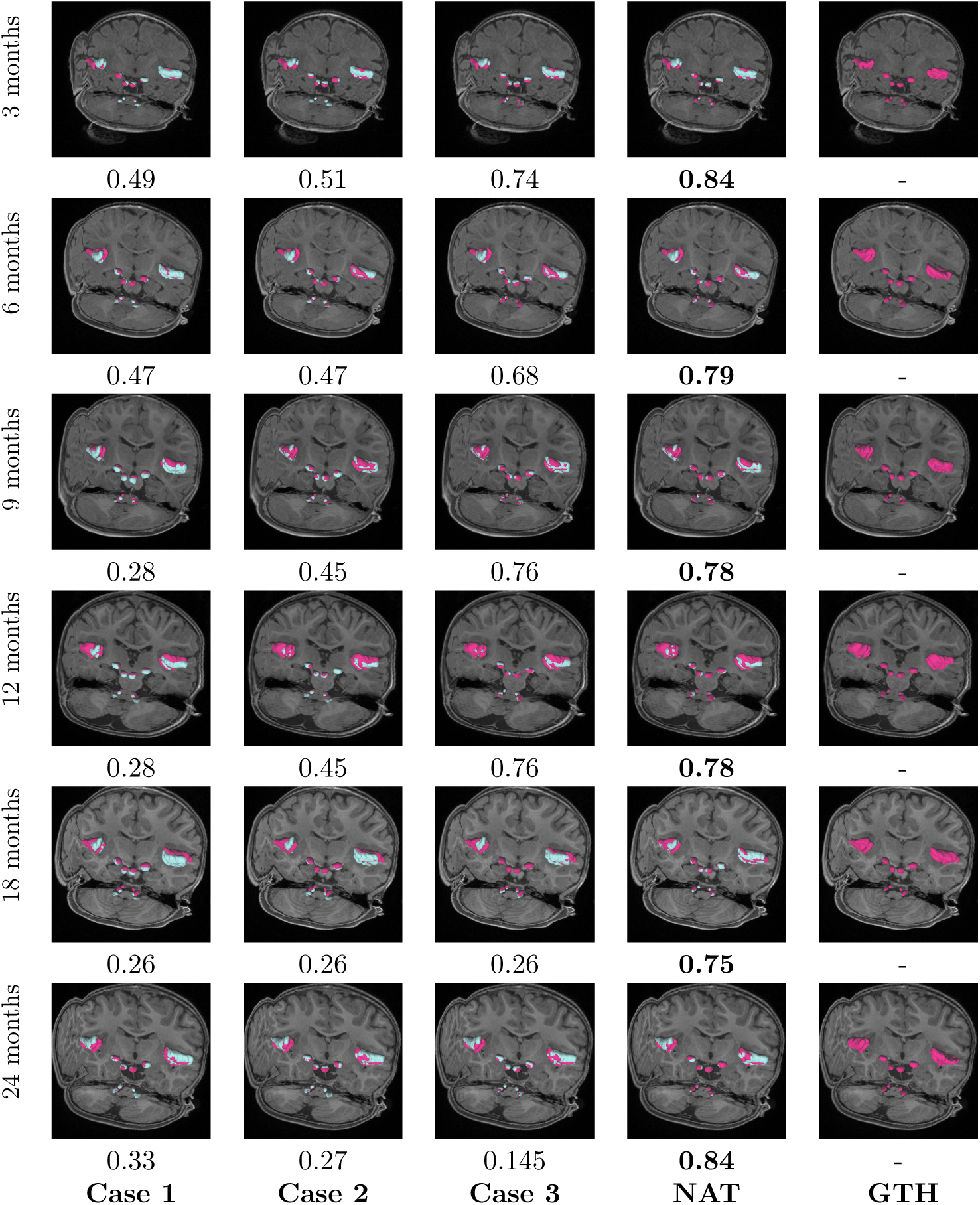
Alignment of auditory nuclei and averaged DICE scores () across four registration strategies: Case 1 - sMRI only; Case 2 - sMRI with tissue maps; Case 3 - sMRI with tissue maps and brainstem masks; and Case 4 - NAT. Incorporating tissue maps enhanced cortical alignment, while the inclusion of brainstem masks reduced errors in auditory nuclei alignment but introduced new alignment issues. The NAT achieved superior performance by separately aligning cortical and brainstem regions.

- **Case** 1: Registration using only sMRI data (T1w and T2w).
- **Case** 2: Registration using sMRI data combined with tissue maps to assist the alignment process.
- **Case** 3: Registration incorporating brainstem masks, in addition to tissue maps, to enhance brainstem alignment.

NAT builds upon a two-branch registration framework that separately aligns cortical and brainstem regions using sMRI data, tissue maps, and brainstem masks (see Fig. 10 in the Method section). As illustrated in Fig. 3, this approach outperforms Cases 1, 2, and 3, with NAT-aligned nuclei consistently exhibiting higher accuracy and closer correspondence to the ground truth (GTH) across all age periods. The GTH nuclei were manually labeled by a neuroradiologist with 15 years of expertise and subsequently reviewed by another with 29 years of experience, ensuring high accuracy and reliability. These findings underscore the robustness of NAT in aligning auditory nuclei under varying developmental periods.

### 2.2 Maturation of auditory tracts

To delineate the developmental trajectories of the auditory system, we characterized the maturation of auditory tracts and examined their correlations with MSEL *t*-scores in four domains: *visual reception*, *fine-motor skills*, *receptive language*, and *expressive language*. We assessed white matter microstructure using two complementary models: neurite orientation dispersion and density imaging (NODDI) [87] and diffusion tensor imaging (DTI) [88]. Specifically, these models were applied to generate microstructural attribute maps, including the neurite density index (NDI), radial diffusivity (RD), and fractional anisotropy (FA), thereby providing detailed information on white matter development throughout infancy.

We computed averaged NDI, RD, and FA values for each auditory tract, then modeled the population trajectories using a linear mixed effect model (LMM) [89]. As reported in Table 2, a quadratic model in the LMM provided superior performance, based on adjusted *r*-squared and Akaike information criterion (AIC), compared to the commonly used linear model. Although linear models are frequently employed to represent early white matter development [78, 90], our results indicate that a quadratic model more accurately captures the non-linear trajectories of auditory tracts from birth to 24 months.

**Table 2.**
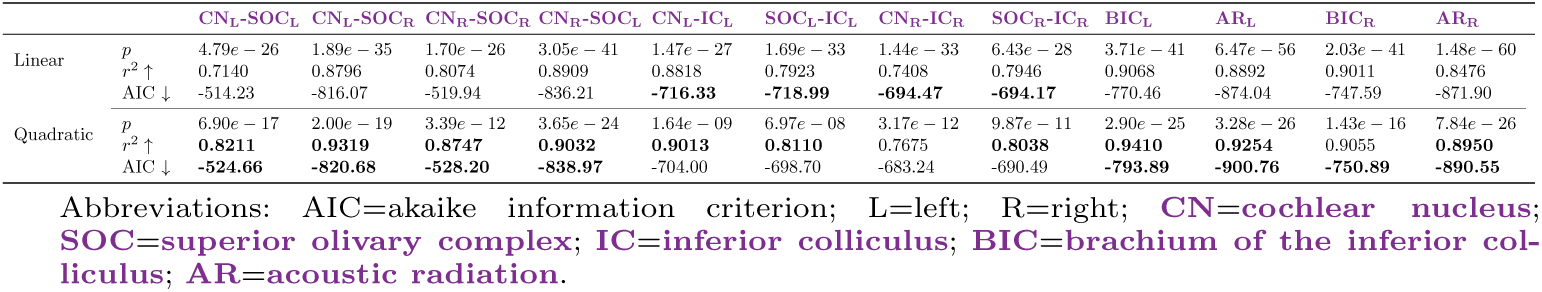
Comparison of linear and quadratic Linear Mixed-Effect Models for auditory tract NDI trajectories based on adjusted *r*-squared and AIC scores.

|  |  | CN<sub>L</sub>-SOC<sub>L</sub> | CN<sub>L</sub>-SOC<sub>R</sub> | CN<sub>R</sub>-SOC<sub>R</sub> | CN<sub>R</sub>-SOC<sub>L</sub> | CN<sub>L</sub>-IC<sub>L</sub> | SOC<sub>L</sub>-IC<sub>L</sub> | CN<sub>R</sub>-IC<sub>R</sub> | SOC<sub>R</sub>-IC<sub>R</sub> | BIC<sub>L</sub> | AR<sub>L</sub> | BIC<sub>R</sub> | AR<sub>R</sub> |
| --- | --- | --- | --- | --- | --- | --- | --- | --- | --- | --- | --- | --- | --- |
| Linear | $p$ | 4.79e-26 | 1.89e-35 | 1.70e-26 | 3.05e-41 | 1.47e-27 | 1.69e-33 | 1.44e-33 | 6.43e-28 | 3.71e-41 | 6.47e-56 | 2.03e-41 | 1.48e-60 |
| | $r^2 \uparrow$ | 0.7140 | 0.8796 | 0.8074 | 0.8909 | 0.8818 | 0.7923 | 0.7408 | 0.7946 | 0.9068 | 0.8892 | 0.9011 | 0.8476 |
| | AIC ↓ | -514.23 | -816.07 | -519.94 | -836.21 | **-716.33** | **-718.99** | **-694.47** | **-694.17** | -770.46 | -874.04 | -747.59 | -871.90 |
| Quadratic | $p$ | 6.90e-17 | 2.00e-19 | 3.39e-12 | 3.65e-24 | 1.64e-09 | 6.97e-08 | 3.17e-12 | 9.87e-11 | 2.90e-25 | 3.28e-26 | 1.43e-16 | 7.84e-26 |
| | $r^2 \uparrow$ | **0.8211** | **0.9319** | **0.8747** | **0.9032** | **0.9013** | **0.8110** | 0.7675 | **0.8038** | **0.9410** | **0.9254** | 0.9055 | **0.8950** |
| | AIC ↓ | **-524.66** | **-820.68** | **-528.20** | **-838.97** | -704.00 | -698.70 | -683.24 | -690.49 | **-793.89** | **-900.76** | **-750.89** | **-890.55** |

Abbreviations: AIC=akaike information criterion; L=left; R=right; **CN=cochlear nucleus**; **SOC=superior olivary complex**; **IC=inferior colliculus**; **BIC=brachium of the inferior colliculus**; **AR=acoustic radiation**.

To facilitate understanding of the extracted auditory tracts, we individually illustrated them in sub-figures within Fig. 4. These panels were arranged from peripheral to cortical regions in a top-to-bottom, left-to-right sequence. For consistency, the same layout applied to Figs. 5, 6, and 7, which depict the developmental trajectories of NDI, RD, and FA, respectively. Note that the CO-CN tracts were excluded from our analysis owing to significant noise in the peripheral microstructural maps, making reliable data interpretation infeasible.

**Fig. 4.**
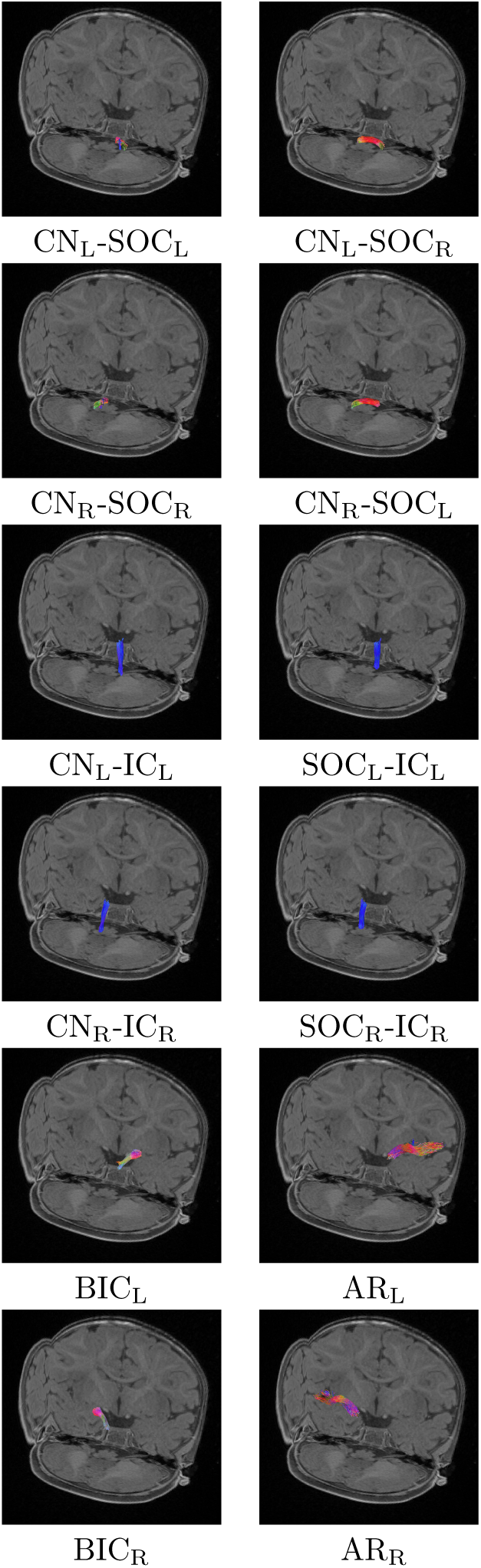
Extracted auditory tracts from dMRI data of a 6-month-old infant. Detailed visualization highlights the small size and complex structure of auditory tracts within the brainstem. (Abbreviations: L=left; R=right; **CN**=**cochlear nucleus**; **SOC**=**superior olivary complex**; **IC**=**inferior colliculus**; **BIC**=**brachium of the inferior colliculus**; **AR**=**acoustic radiation**.)

**Fig. 5.**
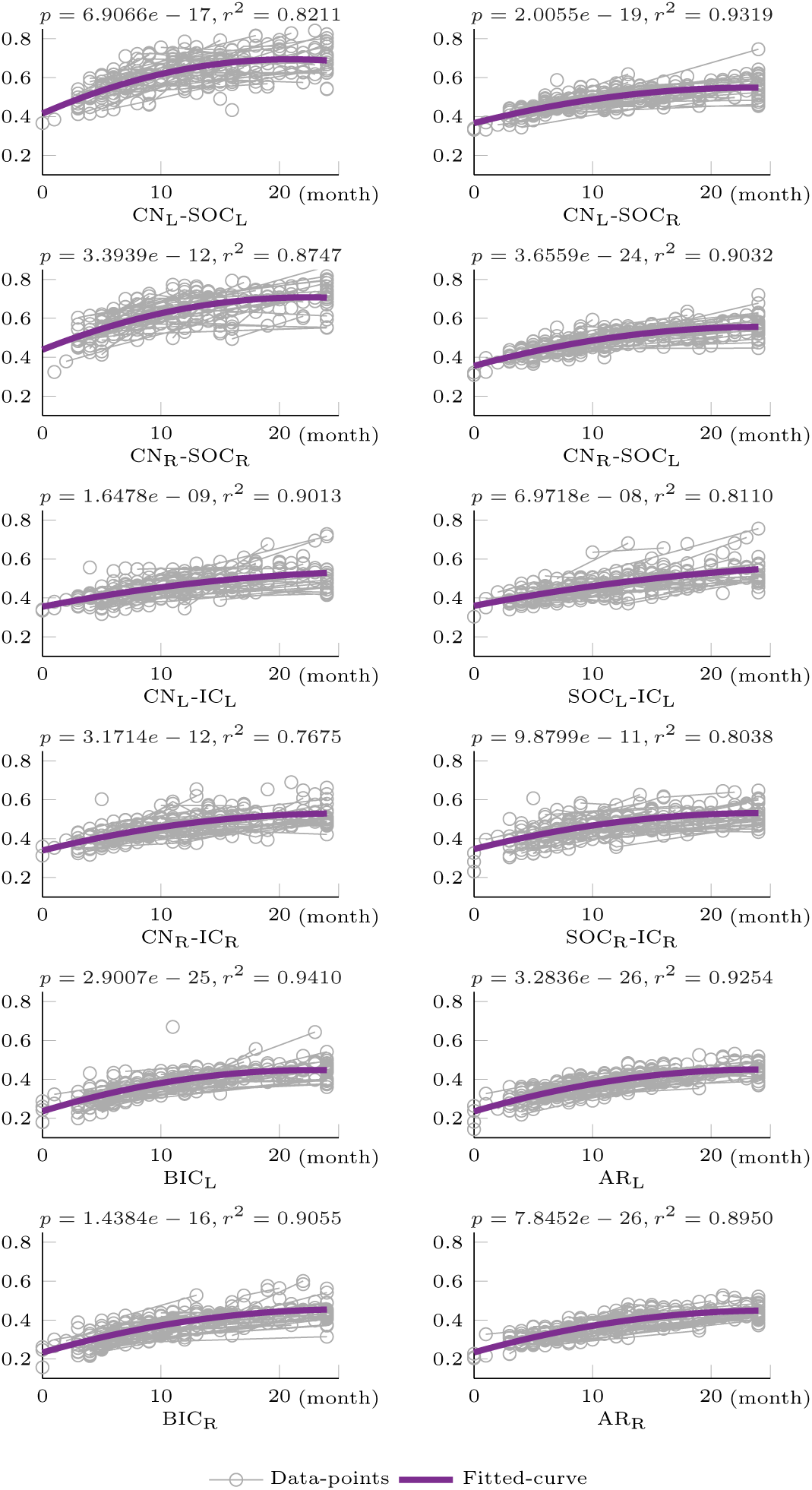
Developmental trajectories of NDI with fitted curves and data points. NDI values gradually increase from birth to 24 months, exhibiting higher neurite density in brainstem regions compared to cortical regions across both hemispheres.

**Fig. 6.**
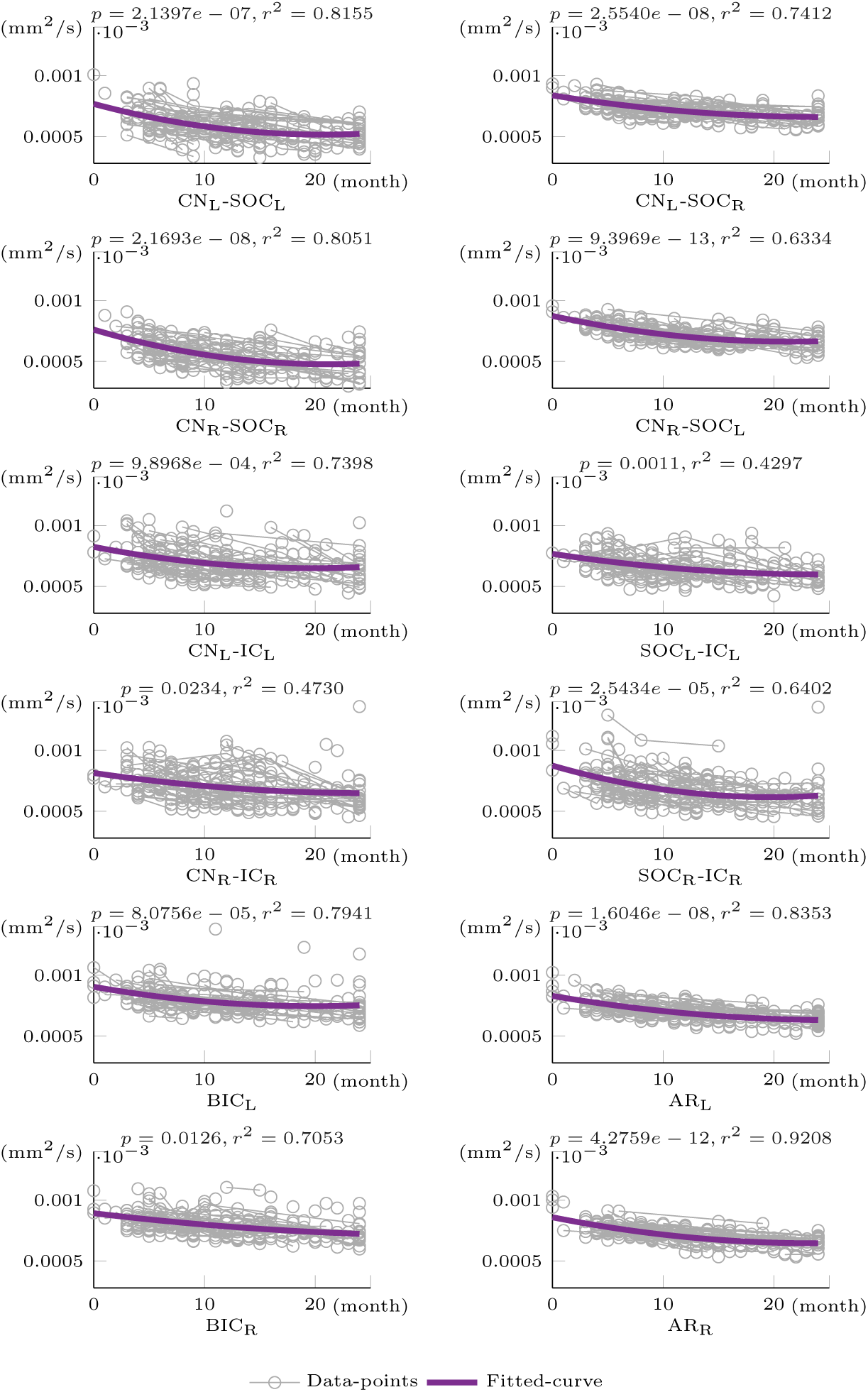
Developmental trajectories of RD with fitted curves and data points. RD values decrease from birth to 24 months in both hemispheres, demonstrating higher myelination near the CN and lower myelination near the PAC.

The NDI ranges from 0 to 1, with values approaching 1 indicating higher neurite density. As illustrated in Fig. 5, the fitted NDI curves for all auditory tracts display a steady increase from birth to 24 months. At birth, tracts situated in brainstem regions exhibit higher maturity levels than those in cortical regions, along the peripheral-to-cortical axis. This pattern aligns with previously proposed developmental progressions of white matter, which follow the direction of information flow [91]. Notably, the developmental trajectories reveal marked asynchrony, with tracts closer to peripheral regions maturing earlier than those nearer to PAC. This finding underscores the staggered timeline of auditory system maturation across infancy.

The inverse of RD reflects myelination status, such that lower RD values indicate better myelination. As illustrated in Fig. 6, RD trajectories gradually decrease over time, closely mirroring the patterns observed in NDI. RD, similar to NDI, shows better development in tracts closer to CN compared to those near the PAC.

The DTI model underlying RD, however, exhibits notable limitations, especially its inability to characterize crossing fibers [68, 92–95]. Consequently, RD-based assessments may be less accurate than NDI (derived from the NODDI model). This limitation becomes more pronounced when using FA, which generally struggles to characterize white matter microstructure in brainstem regions rich in crossing fibers. As shown in Fig. 7, the contralateral CN-SOC exhibits markedly lower FA values compared to the acoustic radiation (AR) and ipsilateral CN-SOC. These FA trajectories do not align with their NDI or RD counterparts, underscoring the challenges in using FA to represent microstructural changes in regions with complex fiber configurations. The NDI values in auditory tracts exhibited significant correlations with fine-motor skills and expressive language *t*-scores from the MSEL tests (see Tables 3 and 4). We used a LMM with a quadratic fit, adjusting all *p*-values for multiple comparisons across tracts via the Benjamini-Hochberg procedure [96].

**Fig. 7.**
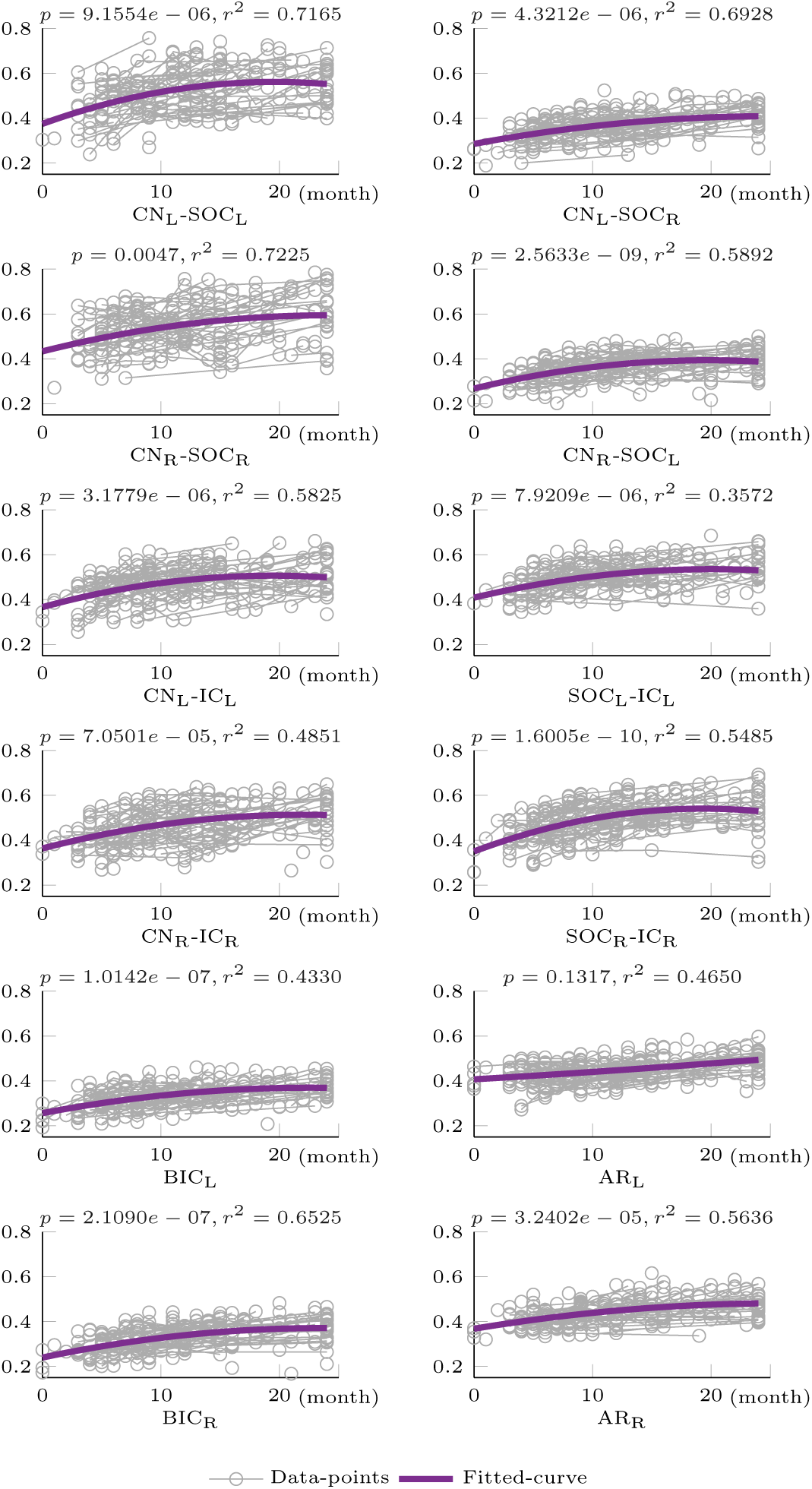
Developmental trajectories of FA with fitted curves and data points. The trajectories in both hemispheres show lower FA in the contralateral CN-SOC compared to the AR and ipsilateral CN-SOC. These patterns reflect the limitations of FA in accurately characterizing white matter microstructure in regions with crossing fibers.

**Table 3.** Significant correlations between auditory tract NDI values and fine-motor skills *t*-scores from the MSEL tests.

| Fine motor $t$-scores | $p$-value | $r^2$ |
| --- | --- | --- |
| $AR_L^*$ | 0.019 | 0.047 |
| $AR_R^*$ | 0.029 | 0.091 |
| $CN_R-IC_R^*$ | 0.025 | 0.050 |
| $CN_R-SOC_R^*$ | 0.015 | 0.135 |

**Table 4.** Significant correlations between auditory tract NDI values and expressive language *t*-scores from the MSEL tests.

| Expressive language $t$-scores | $p$-value | $r^2$ |
| --- | --- | --- |
| $BIC_L^{**}$ | 0.006 | 0.349 |
| $BIC_R^{**}$ | 0.008 | 0.402 |
| $CN_L-IC_L^*$ | 0.013 | 0.333 |
| $CN_R-IC_L^*$ | 0.013 | 0.347 |

For fine-motor *t*-scores, statistically significant correlations (*p <* 0.05) emerged with several auditory tracts, including the bilateral AR, CN_R_-IC_R_, and CN_R_-SOC_R_. Nevertheless, these models showed relatively low *r*-squared scores (below 0.15), suggesting limited explanatory power. This finding implies that, while the auditory tracts do appear to support fine-motor skill development, other systems, such as motor or sensorimotor, may play a more prominent role. Further investigation is needed to clarify these relationships and underlying mechanisms.

In contrast, highly significant correlations were found between NDI values of auditory tracts and expressive language *t*-scores, specifically for the bilateral brachium of the inferior colliculus (BIC) (*p <* 0.01). Additionally, significant correlations were observed with the CN_L_-IC_L_ and CN_R_-IC_L_ tracts (*p <* 0.05), although both tracts did not reach the same level of statistical significance as the BIC. All four tracts exhibited higher *r*-squared scores (*r*-squared*>* 0.3), indicating a moderate explanatory power of auditory systems for expressive language abilities.

These findings underscore the significant contribution of the auditory system to expressive language. The development of expressive language relies on the fine-motor control of speech-specific articulators, such as the tongue, lips, and vocal cords, indicating that the auditory system’s influence is closely linked to these motor functions. Nevertheless, the roles of visual, motor, somatosensory, and sensorimotor systems in supporting speech production should not be overlooked, as they also play crucial roles in this complex cognitive domain. Further research is needed to elucidate the interactions and underlying mechanisms among these systems in the development of language skills.

## 3 Discussion

This study presents a novel toolbox for extracting auditory tracts in infants, providing a robust method to characterize white matter microstructural changes during development. By modeling the gradual and continuous maturation trajectories of these auditory tracts from birth to 24 months, we identified significant correlations between auditory tract NDI values and both *fine-motor skills* and *expressive language t*-scores from the MSEL tests. Notably, these findings offer valuable insights into the intertwined development of sensory and motor systems underlying speech acquisition, particularly in the context of perceptual reorganization, and highlight the importance of investigating how early white matter maturation may contribute to emerging speech abilities.

### 3.1 A robust framework for auditory tract extraction

Research on infant neuroimaging has long faced challenges in data acquisition, processing, and analysis [69, 97]. Despite continuous advancements in data collection and the development of numerous processing and analysis tools [68, 79, 80, 98–106], extracting auditory tracts in infants remains particularly difficult. In previous studies, such extraction often required specimen scanning or ultra-high-field scanners [74–76], and was limited to only a few of the 14 auditory tracts.

To address these limitations, we introduced *NeoAudi Tract* (NAT), a framework specifically developed to extract the full set of auditory tracts, spanning peripheral, subcortical, and cortical regions, from 3T HARDI data. NAT was designed to accommodate significant variability in infant MRI data, with a core emphasis on maximizing registration robustness. The key contributions of the NAT framework, which enable robust auditory tract extraction, include: (1) An auditory tract atlas constructed from the Human Connectome Project (HCP) dataset [107], offering high-quality anatomical references; (2) The use of high-spatial- and high-temporal-resolution UNC 4D infant templates [103], to preliminarily normalize brain orientation and field of view (FOV); and (3) The integration of tissue and brainstem segmentation maps, which provide additional anatomical constraints to accurately align cortical and brainstem regions in separate registration branches. Because prior processing steps often retain neck structures in the FOV [99], a two-branch registration framework is essential for handling the resulting variability in neck size, ensuring accurate alignment.

Collectively, these design elements enable NAT to achieve accurate alignment of both cortical and brainstem nuclei across different developmental periods, thereby facilitating the extraction of the full set of auditory tracts. Using these extracted tracts, we observed a progressive increase in tract size and streamline density, reflecting a gradual and continuous developmental course from birth to 24 months. This observation is further supported by the modeled developmental trajectories, which provide quantitative evidence of ongoing white matter maturation throughout early infancy.

The NDI, RD, and FA trajectories for each tract exhibit steady increases or decreases over the 24-month period. Noticeably, tracts closer to peripheral regions exhibit higher maturity compared to those near the PAC, as indicated by the NDI and RD trajectories. However, FA values reveal pronounced differences in maturity levels between peripheral and cortical tracts. This discrepancy likely arises from the limitations of the DTI model in representing crossing fibers [68, 92–95], especially within the brainstem regions where fiber crossing is more prevalent. These findings underscore the importance of using high-quality imaging data, such as HARDI, and advanced diffusion models like NODDI. By offering more reliable microstructural information, these techniques can more accurately characterize the underlying neural substrate changes, thereby refining our understanding of early neurodevelopment.

Despite its overall success, NAT has certain limitations, as reflected in an average DICE score of approximately 0.8 for nuclei alignment. In particular, the low signal-to-noise ratio (SNR) in peripheral regions, coupled with the structural complexity of the brainstem, poses challenges for precisely extracting small tracts such as the cochlear nerve and CN–SOC pathways. Future improvements in dMRI acquisition, including higher spatial and angular resolution, as well as advancements in extraction methods, will be critical to overcoming these difficulties. Additionally, constructing age-specific auditory tract atlases would further enhance extraction accuracy and reliability by better accounting for developmental variations across infancy.

### 3.2 A potential role of the auditory system in speech production

The auditory system supports the perception of acoustic stimuli by converting speech vibrations into electrical signals and enabling subsequent analyses of spectral, intensity, temporal, and spatial features [82, 108]. However, speech inherently involves both acoustic and articulatory dimensions [109, 110], with activity coordination in sensory and motor systems [65]. Recent research has therefore examined not only how the auditory or motor system independently influences speech perception [22, 32], but also how the sensorimotor system integrates these processes [20, 45]. Investigators emphasize that both sensory and motor experiences are essential for effective learning [53–55].

This raises a critical question: How do the auditory and motor systems develop in tandem to support speech acquisition, especially in the context of perceptual reorganization? While this question is fundamental to understanding the developmental relationship between sensory and motor systems for functional integration, it remains unresolved.

Current exploration of this topic still faces challenges in characterizing those tiny and complex nuclei structures in peripheral and brainstem regions [68]. To investigate the role of the auditory system in speech acquisition, we developed NAT and examined correlations between auditory tract maturation and four domains of the MSEL: visual reception, fine-motor skills, receptive language, and expressive language. Among these domains, the NDI values of auditory tracts showed statistically significant correlations with *t*-scores in fine-motor skills and expressive language, suggesting a link between auditory processing and speech production. Our findings, along with those highlighting the influence of the motor system on speech perception [20, 45], raise the possibility that the development of speech perception and production is intertwined, pointing to further research into the integrated development of sensory and motor systems.

Despite the repeatedly reported link between speech perception and production in infants [18, 35, 42, 44], *auditory–motor associations may not emerge innately* [48]. Instead, recent research suggests that postnatal sensorimotor experiences facilitate the establishment of these associations [111, 112], forming a basis for imitation [50, 52, 113]. However, the origin of imitation remains a topic of heated debates [114–127].

Once these sensory–motor associations are formed, sensory input can simultaneously activate motor regions [128, 129], potentially driving plastic changes in both auditory and motor networks underlying speech acquisition [57]. During the refinement of neural connectivity, the auditory and motor systems gradually become more attuned, contributing to the sharpening of infants’ sensitivity to their native language. One more specific hypothesis is that the auditory system may influence perceptual reorganization via mediation by the motor system. However, further investigation is needed to clarify the precise mechanisms underlying how auditory and motor system development interact to support early speech activities.

Among these auditory tracts, we found no significant correlation with receptive language *t*-scores from the MSEL tests. One possible explanation may lie in the emphasis on semantic comprehension in the test items, which are more likely to engage ventral language pathways that map sound to meaning [61, 110, 130]. Consequently, these semantic skills could be less dependent on the specific auditory tracts examined in our study.

In light of this, future research would expand NAT’s focus beyond the auditory system to include additional systems, such as sensorimotor, visual, and motor tracts [131, 132], to gain a more comprehensive understanding of how interconnected neural networks support speech acquisition and cognitive development during early infancy. Such an integrative approach may reveal how ventral and dorsal language pathways, along with additional sensory and motor pathways, collectively shape both receptive and expressive language outcomes.

#### Concluding remarks

We propose a novel toolbox, *NeoAudi Tract* (NAT), for extracting auditory tracts in infants. By charting their maturation trajectories, we identified statistically significant correlations between the NDI values of these tracts and *t*-scores for *fine-motor skills* and *expressive language* from the MSEL tests, thereby supporting the linked development of speech perception and production. We hypothesize that speech perception and production are mutually dependent processes, driven by the intertwined maturation of sensory and motor systems.

Beyond advancing our understanding of typical auditory tract maturation, NAT also holds promise for investigating developmental deviations in hearing-impaired infants. Studies have demonstrated that early auditory implants can significantly improve speech performance and language-learning outcomes [133–136], although the optimal timing and implantation strategy remain to be fully established. By uncovering both typical and atypical maturation trajectories in hearing-impaired and congenitally deaf infants [137–139], NAT can provide critical insights to inform clinical decision-making and predict postoperative outcomes.

In line with the paradigm shift from studying isolated brain regions to examining interconnected networks [58, 61], cognitive functions are increasingly regarded as emergent properties of brain-wide interactions [58, 59, 140]. This perspective underscores the need to investigate how specific neural systems, such as the auditory and motor systems, develop and interact to support behavioral outcomes [132]. Our results indicate that the maturation of auditory tracts is closely associated with speech production within the context of motor system development, providing valuable insights into the plastic changes in white matter that integrate functionally segregated regions and underpin cognitive behaviors. Furthermore, this work highlights the potential of task-free, longitudinally applicable dMRI data from a 3T scanner — capable of simultaneously characterizing multiple systems — to advance our understanding of speech acquisition mechanisms. Looking ahead, future investigation will benefit from interdisciplinary collaboration across fields such as linguistics, neuroscience, psychology, neuroimaging, medicine, and computer science [2]. Such efforts will ultimately deepen our knowledge of how neural substrates develop to subserve early language and cognitive development.

## 4 Method

To reliably extract auditory tracts from infant dMRI data, we first constructed a high-quality atlas of auditory tracts using HCP adult data [107], then employed our proposed toolbox for processing infant sMRI and dMRI data [99], and finally designed a minimal processing pipeline to ensure accurate registration when aligning adult templates with infant data. The following sections will introduce the main framework of NAT (Section 4.1), the procedures for constructing the adult auditory tract atlas (Section 4.2), and the minimal processing steps for infant sMRI (Section 4.3) and dMRI data (Section 4.4).

### 4.1 NAT framework

As illustrated in Fig. 8, NAT comprises two main components: (1) *atlas preparation* and (2) *tract extraction*. NAT is specifically designed to improve registration accuracy, ensuring reliable alignment between infant data and adult templates. By simply replacing the atlas and segmented ROI masks, NAT can be easily adapted to extract any tract in subjects at any age period, offering a flexible solution for a broad range of developmental neuroimaging studies.

**Fig. 8.**
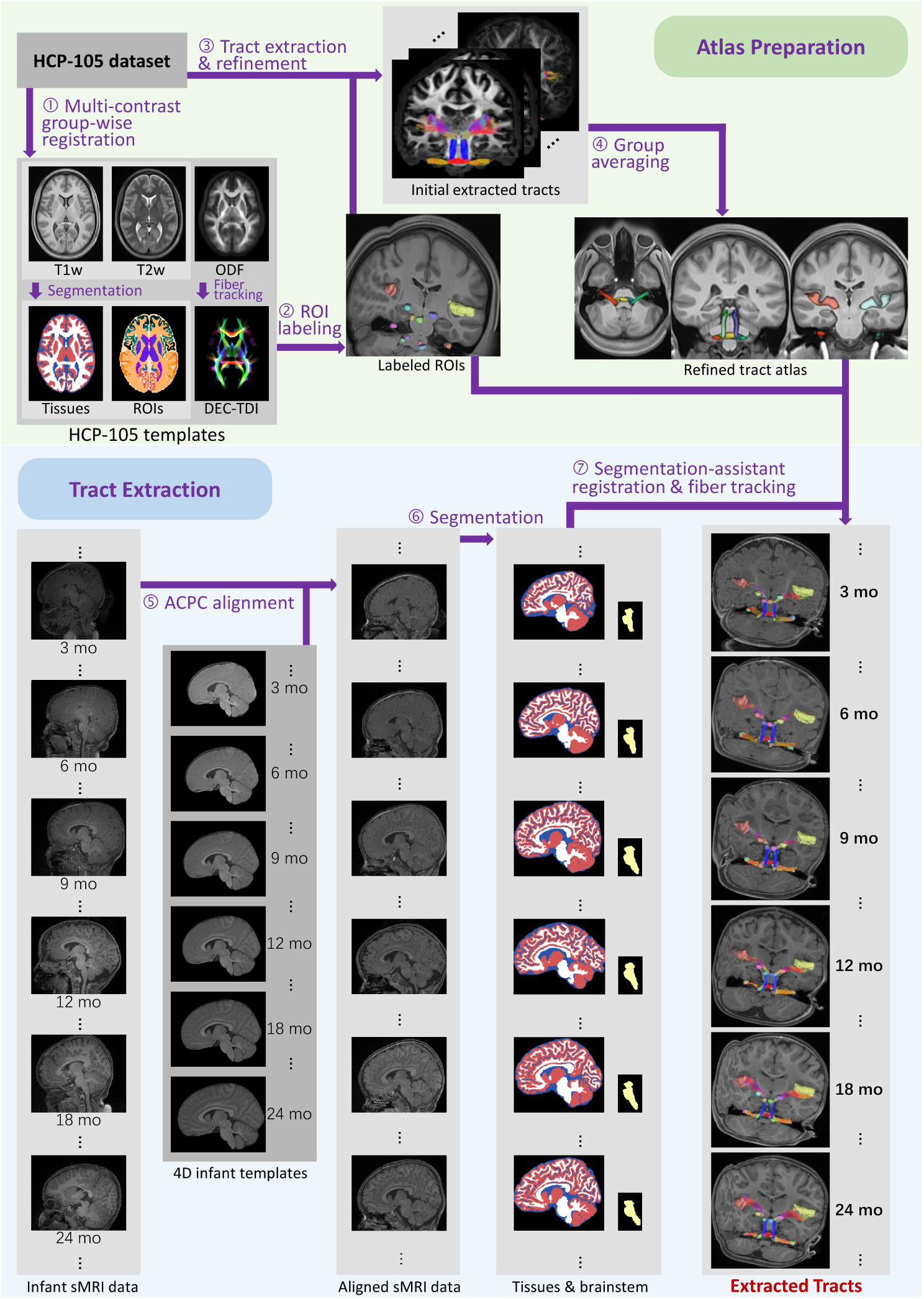
Overview of the NAT framework, which comprises two main components: (1) *atlas preparation* and (2) *tract extraction*. (1) Atlas preparation involves creating a high-resolution atlas of auditory tracts, including key nuclei such as the cochlea (CO), cochlear nucleus (CN), superior olivary complex (SOC), inferior colliculus (IC), medial geniculate body (MGB), and primary auditory cortex (PAC; Heschl’s gyrus), as well as tract masks. (2) Tract extraction utilizes precise segmentation of infant brain tissues and ROIs to support the registration framework, enabling accurate delineation of tract anatomy.

In the first part, *atlas preparation*, we employed a high-quality MRI dataset from HCP [107], which enabled us to delineate the minute nuclei in auditory pathways. Below, we detail four core procedures: **(1.1)** We first constructed group-wise, multi-modality MRI templates from 105 subjects [141]. **(1.2)** Using complementary information revealed in these multi-modality MRI templates, one experienced neuroradiologist (15 years of experience) labeled the initial and terminal regions of the auditory tracts. Another neuroradiologist (29 years of experience) then reviewed these labels to ensure accuracy. **(1.3)** The labeled ROIs in the template space were aligned to each subject’s space to facilitate the extraction of auditory tracts. **(1.4)** The extracted tracts in each subject space were subsequently aligned back to the template space and group-averaged, resulting in an initial tract atlas. We used this initial tract atlas to refine the tract extraction in Step (1.3), and the newly extracted tracts were employed to update the tract atlas in Step (1.4). These two steps were repeated iteratively, thereby refining the tract atlas with each iteration.

In the second part of NAT, *tract extraction*, we performed three core procedures:

**(2.1)** We first aligned the infant sMRI data with the UNC 4D infant templates for anterior commissure and posterior commissure (ACPC) alignment [103], which facilitates subsequent steps, including tissue segmentation and registration. **(2.2)** We then segmented infant tissues and ROI maps, which were incorporated into the registration to improve robustness. **(2.3)** Finally, we combined anatomical information from both the aligned atlas and segmentation maps to ensure the reliability of the extracted auditory tracts.

### 4.2 Atlas preparation and refinement

Given that auditory nuclei are very small and structurally complex, neuroradiologists have to locate them using high-quality T1-weighted (T1w), T2-weighted (T2w), and dMRI data to fully exploit their complementary information [68]. Therefore, we employed 105 subjects from the HCP dataset to construct multi-modality templates for labeling these nuclei efficiently [107, 141]. Instead of using raw data, we employed the processed sMRI and dMRI data provided by HCP [142], which required only a few additional processing procedures as detailed below. It is worth noting that we directly employed the 0.7 mm^3^ T1w and T2w data, rather than the 1.25 mm^3^ data accompanying the processed dMRI data, to preserve higher spatial resolution information. We aligned the T1w and T2w data during ACPC alignment, ensuring consistent orientation across modalities.

As shown in Fig. 9, we constructed both the sMRI and dMRI (*i.e.*, fiber orientation distribution function (ODF)) templates for labeling auditory nuclei. To build T1w and T2w templates, we employed the *antsMultivariateTemplateConstruction2* script from the ANTs toolbox [143, 144], which iteratively generates and optimizes candidate templates by incorporating multi-modality information [145]. Before constructing the ODF template, we modified our previously proposed dMRI processing pipeline [99] to better accommodate peripheral and brainstem regions. Specifically, we first estimated the response function for spherical deconvolution using the *dhollander* algorithm [146], then chose the *csd* approach for generating constrained spherical deconvolution ODFs [147]. To build ODF template, we employed the *population template* tool from MRTrix3 [148]. In order to properly include peripheral tracts, we dilated the brain masks before normalizing ODFs, thereby ensuring these regions were not inadvertently excluded.

**Fig. 9.**
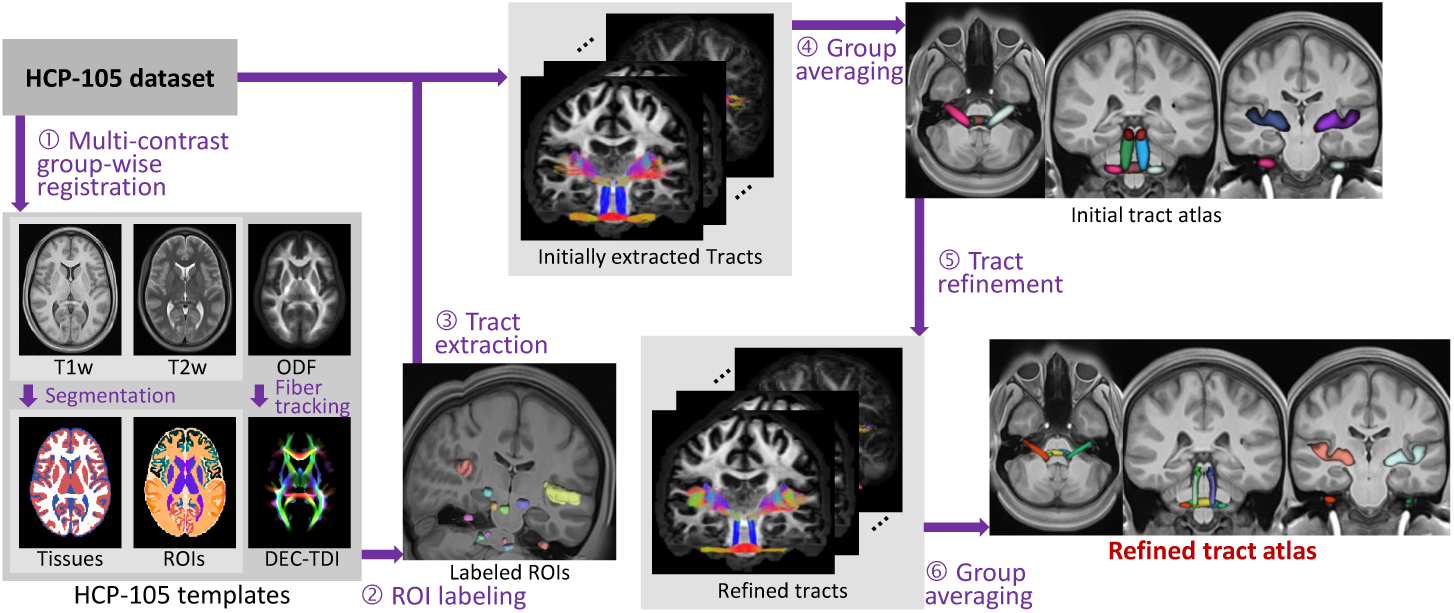
Preparation of high-resolution templates and auditory tract atlas for adults. Labeled ROIs in the template space were used to extract initial tracts, which were then utilized to create tract probability maps, forming a preliminary tracts atlas. This atlas was subsequently refined to enhance the precision of auditory tract extraction, resulting in a more accurate tracts atlas.

Once the T1w, T2w, and ODF templates were constructed, we aligned the T1w template to the ODF template using ANTs rigid registration [149]. We then applied deep-learning–based segmentation methods to the T1w template in order to obtain tissue maps and ROIs (including Desikan–Killiany–Tourville (DKT) regions as well as subcortical and cerebellar areas) [150, 151]. Next, the T2w template, along with tissue and ROI maps, was warped to the dMRI data space using the same transformation matrix derived from the T1w-template registration. In addition, we used the ODF template to generate a directionally-encoded color track density imaging (DEC-TDI) map, following the configurations described in [68]. Finally, experienced neuroradiologists delineated the auditory nuclei by jointly examining the T1w, T2w, DEC-TDI, tissues, and DKT-ROI maps.

The labeled auditory ROIs were then warped to each subject’s dMRI space, serving as the initial and terminal regions for fiber tracking. These aligned ROIs enabled the extraction of initial auditory tracts in the individual dMRI space. All extracted tracts were subsequently aligned back to the template space for group averaging, resulting in an initial tract atlas. Because the initial auditory tracts might include outlier streamlines, we further used the initial auditory tract atlas to discard streamlines lying outside the aligned atlas. These refinement procedures can be repeated iteratively; in this work, we refined the initial atlas only once, as it already demonstrated the adequate accuracy.

### 4.3 Minimal processing for sMRI data

During the first two postnatal years, the infant brain undergoes rapid growth in size and ongoing myelination, leading to substantial changes in gray matter (GM) and white matter (WM). Consequently, sMRI data of this period exhibit large variations in intensity distributions [106], posing significant challenges for accurate brain tissue segmentation and registration. Hence, robust sMRI data processing is crucial for leveraging the adult atlas to extract auditory tracts from infant neuroimaging data. Building on our previously proposed tools for infant neuroimaging processing [68, 98, 101, 104], we obtained brain tissue and ROI maps from neonates up to 24-month-old infants. This minimal processing framework helps ensure reliable alignment between infant sMRI data and the adult templates, forming the foundation for subsequent auditory tract extraction.

#### 4.3.1 sMRI data preprocessing

We performed sMRI data preprocessing using FSL [152], ANTs [153], and MRtrix3 [148]. First, all sMRI data underwent N4 bias field correction to address intensity inhomogeneities [153]. We then obtained a rough brain mask by rigidly aligning each T1w data to the corresponding age-specific template [103], which allowed for a rough skull stripping and removal of facial and neck tissues. Next, the skull-stripped T1w data served as a reference for ACPC alignment, so that improving the robustness of subsequent brain tissue segmentation and non-rigid registration. Finally, for each subject, the T2w data were rigidly aligned to the T1w data using *flirt* in FSL, ensuring consistent orientation across both modalities.

Subsequently, we employed our previously proposed deep learning models to segment infant brain tissues [101, 104]. These models leverage the intensity-invariant anatomical features to accommodate substantial variations in intensity distributions characteristic of the rapid development. Unlike prior studies that relied on age-specific models [154], our single-model approach enabled precise and longitudinally consistent tissue segmentation across a broad age range [104]. Both tissue and brainstem segmentation maps then served as anatomical guidance for accurately aligning infant sMRI data with adult templates, ensuring robust registration despite the considerable variations in intensity distribution.

#### 4.3.2 Two-branch registration framework

The auditory tract extraction poses two major challenges for the alignment: (1) auditory tracts traverse peripheral, subcortical, and cortical regions, and (2) the lower portions of the brainstem exhibit large variations in the processed neuroimaging data. Consequently, it is difficult to simultaneously align both the cortical and brainstem regions of adult templates to infant sMRI data. To address this issue, we developed a two-branch registration framework that separately aligns the cortical areas and the brainstem, as illustrated in Fig. 10. This design allows for more accurate alignment of both regions despite their distinct anatomical and developmental characteristics.

**Fig. 10.**
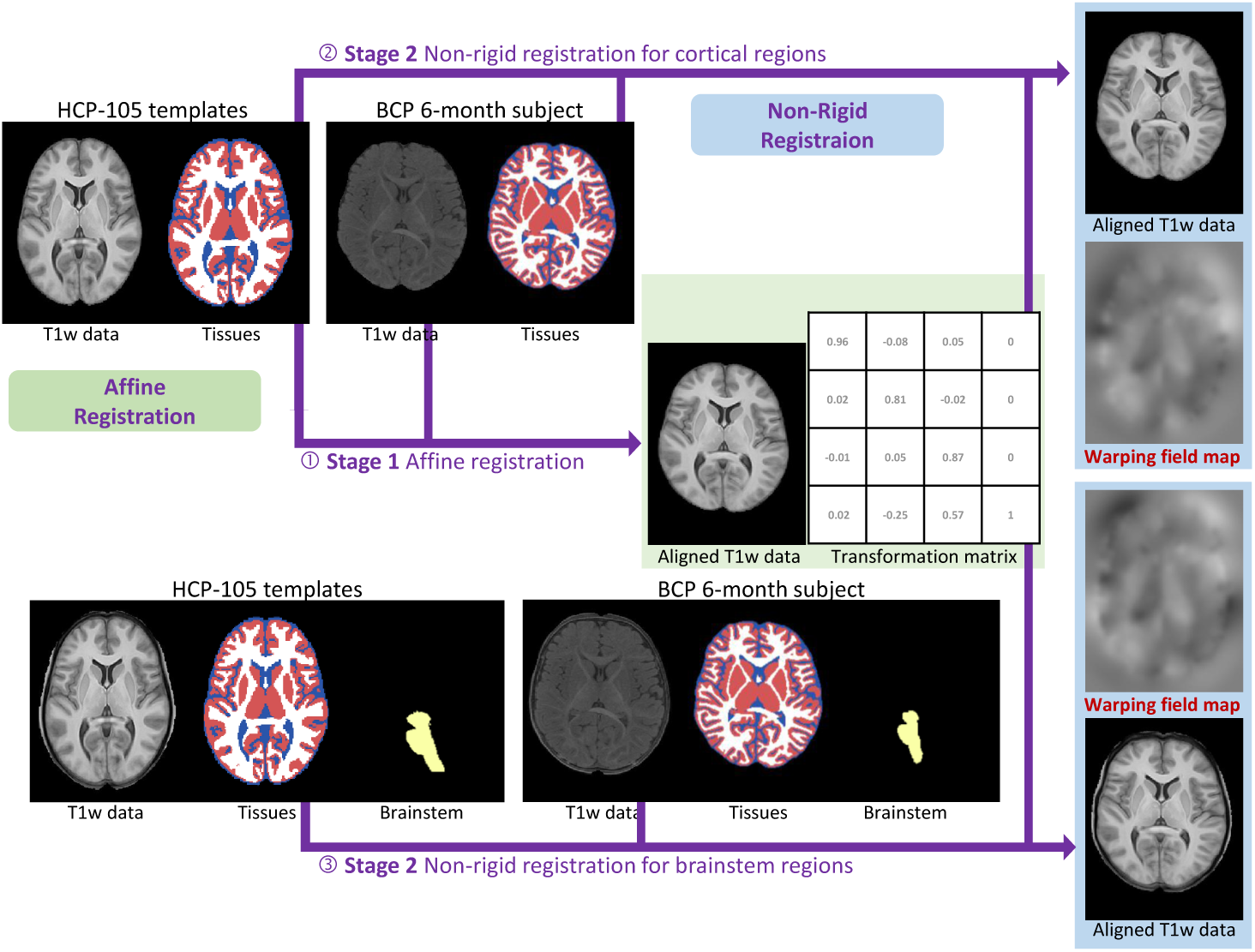
Two-branch registration framework for aligning adult templates with infant sMRI data. The framework utilizes both T1w and T2w data, although T2w data are not shown here for clarity. The alignment process consists of two stages: (1) Affine registration using tissue maps to adjust intensity contrasts between gray matter and white matter; (2) Dual deformable registration branches to finetune the alignment of cortical and subcortical regions separately.

Our proposed framework proceeds in two distinct stages: (1) *affine registration* and (2) *deformable registration*. *In the first stage*, there is a single branch focusing on cortical alignment. We employed skull-stripped data for affine registration of cortical regions (from adult sMRI templates to infant sMRI data), guided by the segmented tissue maps. This process yielded a transformation matrix that serves as the initial transform for the subsequent deformable registrations. *In the second stage*, two branches of deformable registration were performed to separately warp cortical and brainstem regions. For the cortical branch, tissue maps are utilized. For the brainstem branch, in addition to tissue maps, we also incorporated the brainstem map and dilated brain mask to ensure accurate normalization of the brainstem area. Finally, these two-branch warping fields effectively align both cortical and brainstem regions, laying the groundwork for reliable auditory tract extraction.

#### 4.3.3 ROIs refinement for tracking fibers

After completing the registration steps, we obtained reference information from adult atlas. In addition to the aligned auditory ROIs and tract masks in the individual space, we also utilized tract endpoint pairs as complementary cues for refining region definitions. Specifically, we extracted the initial and terminal regions of each tract from its corresponding tract mask. The order of the initial and terminal regions was established, during the construction of the adult tract atlas, by referencing one of the 105 adult subjects using *QuickBundle* [155]. This approach helps ensure a consistent orientation of fibers (*i.e.*, initial *vs*. terminal) across subjects, thereby improving the reliability of subsequent fiber tracking and analysis.

For ROI refinement, we first combined the two endpoint regions of each tract with the corresponding auditory ROIs, then further refined them using segmentation maps, as shown in Fig. 11. This procedure aligns with the anatomically constrained tractography (ACT) framework implemented in MRTrix3 [148]. Specifically, to extract auditory tracts under the ACT framework, we delineated the GM/WM interface and used it to identify intersections between this interface and the combined auditory ROIs. In line with ACT’s objective, these refinement steps help ensure the fidelity of the tracked fibers.

**Fig. 11.**
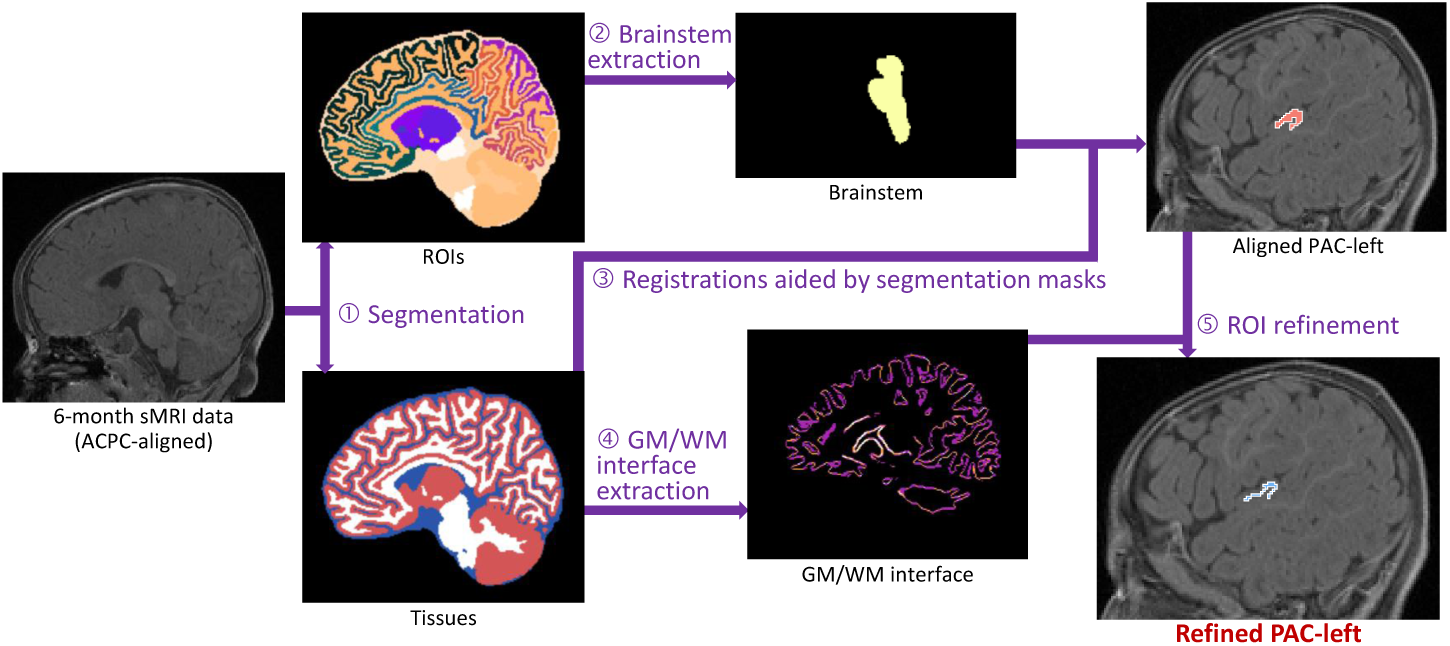
Deep learning based segmentation of brain tissues and DKT-based ROIs using T1w and T2w data. The segmented maps were utilized to delineate the brainstem and the interface between GM and WM. This segmentation provides consistent tissue contrast, facilitating subsequent registration processes, and allows for precise adjustment of aligned ROIs for accurate identification of auditory tracts. The refined left primary auditory cortex (PAC) demonstrates improved accuracy.

### 4.4 Extraction of auditory tracts

To precisely extract auditory tracts, NAT leveraged anatomical guidance from sMRI data to effectively preprocess infant dMRI data [99], thereby alleviating the spatiotemporal asynchrony of white matter development.

We employed MRtrix3, FSL, and ANTs to process the dMRI data. All data underwent bias and distortion correction [156–159], denoising [160], and co-registration [161]. Specifically, we aligned each sMRI data to the dMRI space via boundary-based registration (BBR) [162], guided by the segmented tissue maps. Subsequently, we applied rotation and correction of the B-matrix to ensure accurate streamline tracking [163]. Notably, this differs from the previous pipeline [99] in how the response function and ODF are generated (see Section 4.2 for details), catering to peripheral and brainstem regions. Rather than using a dilated brain mask, we adopted the regular brain mask to normalize the obtained ODF across GM, WM, and cerebrospinal fluid (CSF) [164], thus capturing appropriate tissue information without introducing spurious peripheral influences.

With the normalized ODF, we next utilized the aligned ROIs in the dMRI space to create three types of masks: inclusion, seed, and exclusion. The auditory tracts were then extracted individually using the *iFOD2* algorithm [165]. Specifically, we employed the combined ROIs as inclusion masks, and defined seeds only at regions intersecting the GM/WM interface to ensure biologically plausible in generating streamlines. Meanwhile, we dilated the aligned tract masks and inverted them to derive exclusion masks, thereby filtering out outlier streamlines.

Moreover, we implemented a data quality control (QC) procedure at each step to ensure the reliability of the analyses [166]. Specifically, during QC, each data sample was labeled as *pass*, *questionable*, or *fail*. We excluded *fail* data from subsequent analyses, while both *pass* and *questionable* data were retained.

The QC criteria encompassed three major aspects: (1) acquisition issues (*e.g.*, early stopping, excessive motion, blurring, and signal loss), (2) processing issues (*e.g.*, severe intensity artifacts, distortion-correction failures, misalignment, and tract extraction failures), and (3) tract attributes that were not longitudinally consistent in longitudinal charting. Data labeled *pass* showed no apparent artifacts, good alignment, and consistency across time points, whereas *questionable* datasets exhibited moderate issues in one or more of these areas. Together, *pass* and *questionable* data constituted only 42.89 % of the total dataset.

These configurations ensure that fiber tracking remains confined within valid anatomical boundaries, thereby enhancing the accuracy and consistency of the extracted tracts and ultimately ensuring the reliability of the analysis.

## Declarations

This manuscript is an extended version of the abstract presented at the 2024 *International Society for Magnetic Resonance in Medicine (ISMRM) Annual Meeting*. The abstract was part of the ISMRM 2024 *Power Pitch Program* (#0240). For more details, refer to the following links:

- NAT — ISMRM Abstract Archive Link
- Infant multi-modality MRI data processing — ISMRM Abstract Archive Link
- YouTube Presentation Video

## Acknowledgments

This work was supported in part by the STI 2030-Major Projects (No. 2022ZD0209000), the National Natural Science Foundation of China (No. 62203355, 62073260, 62131015, and U23A20295), the STI 2030-Major Projects (No. 2022ZD0213100), Shanghai Municipal Central Guided Local Science and Technology Development Fund (No. YDZX20233100001001), the Key R&D Program of Guangdong Province, China (No. 2023B0303040001, 2021B0101420006), and HPC Platform of ShanghaiTech University.

## Author Contributions

F.L. and Y.W. designed the research, wrote the codes, and drafted the manuscript. F.L., J.G., and J.L. collected data, designed processing tools, and processed the data. F.L. and X.C. provided statistical analysis and interpretation of the data. Y.W., Z.W., and H.W. made contributions in the areas of anatomical delineation and data analysis. H.Z., J.F., and D.S. coordinated and supervised the whole work. All authors were involved in critical revisions of the manuscript, and have read and approved the final version.

